# ATM-dependent RHEB phosphorylation couples DNA damage to lysosomal mTORC1 signaling to orchestrate the cellular response to genotoxic stress

**DOI:** 10.64898/2026.01.16.699946

**Authors:** Jiyoung Pan, Aurelio A. Teleman, Constantinos Demetriades

**Affiliations:** Max Planck Institute for Biology of Ageing (MPI-AGE), 50931 Cologne, Germany; Cologne Graduate School of Ageing Research (CGA), 50931 Cologne, Germany; German Cancer Research Center (DKFZ), 69120 Heidelberg, Germany; Heidelberg University, 69120 Heidelberg, Germany; University of Cologne, Cologne Excellence Cluster for Aging and Aging-Associated Diseases (CECAD), 50931 Cologne, Germany; European Research Institute for the Biology of Ageing (ERIBA), University of Groningen (RUG), University Medical Center Groningen (UMCG), 9713 GZ Groningen, The Netherlands

**Keywords:** mTORC1 signaling, RHEB GTPase, ATM, TFEB, DNA damage response (DDR)

## Abstract

Cells dynamically adapt to environmental stressors by rewiring signaling networks that coordinate growth, metabolism and genome maintenance. The DNA damage response (DDR) and mTORC1 signaling pathways govern DNA repair and cell growth, respectively, but how these pathways intersect remains incompletely understood. Here, we identify RHEB, the most direct mTORC1 activator, as a substrate of the DDR kinase ATM. Strikingly, we find that, although genotoxic stress differentially regulates mTORC1 activity—reducing the phosphorylation of its lysosomal target TFEB, while enhancing phosphorylation of its cytoplasmic target S6K—the DDR-induced phosphorylation of RHEB specifically controls the lysosomal mTORC1 signaling branch. Preventing RHEB phosphorylation impairs TFEB nuclear translocation and lysosome biogenesis upon DNA damage. Functionally, the RHEB phosphorylation-dependent TFEB response is required for proliferative recovery following genotoxic stress. These findings uncover an ATM-RHEB-mTORC1-TFEB signaling axis that links DNA damage to selective mTORC1 outputs, revealing a mechanism which enables cells to adapt to genotoxic cues.

## Introduction

Cells constantly encounter genotoxic and metabolic stress, and their ability to sense and respond to these cues is essential for maintaining homeostasis and genomic integrity. Two central pathways orchestrating these adaptive responses are the DNA damage response (DDR) and the mechanistic target of rapamycin complex 1 (mTORC1) pathway. While each has been extensively studied, how DNA damage influences mTORC1 activity—and whether these two systems functionally intersect—remains incompletely understood.

The DDR is activated by a broad spectrum of genotoxic stresses arising from both exogenous and endogenous sources. Ionizing radiation, ultraviolet light, and chemical agents represent common environmental challenges, while intracellular processes such as DNA replication and cellular metabolism generate replication stress and reactive oxygen species, respectively, that also compromise genome integrity. These diverse insults converge on a core signaling cascade coordinated by the apical kinases ATM (ataxia-telangiectasia mutated), ATR (ataxia telangiectasia and Rad3 related) and DNA-PK (DNA-dependent protein kinase), which initiate and propagate the cellular response through regulation of cell cycle arrest, DNA repair, senescence and apoptosis^1^.

mTORC1 integrates nutrient, energy and growth factor signals to coordinate anabolic and catabolic processes, including protein synthesis, lysosome biogenesis, autophagy, and protein secretion^2–9^. Its activity is regulated by distinct sets of small GTPases: the heterodimeric Rag GTPases (hereafter, the Rags) mediate mTORC1 recruitment to lysosomes, while the small GTPase RHEB directly binds and activates the complex^10–15^. The Rags function as obligate heterodimers, with RagA or RagB pairing with RagC or RagD^10–13^. Under nutrient-rich conditions, the Rag heterodimer adopts an active conformation, in which RagA/B is GTP-loaded and RagC/D is GDP-loaded. mTORC1 phosphorylates multiple downstream effectors, including its canonical substrates such as S6K (S6 kinase) and 4E-BP1 (eukaryotic translation initiation factor 4E-binding protein 1), as well as the lysosome-associated transcription factors TFEB (transcription factor EB) and TFE3 (transcription factor E3)^16–20^. While RHEB activity is classically controlled by its GTP/GDP loading status, regulated by the TSC (tuberous sclerosis complex) protein complex^21^, recent studies have shown that RHEB is also subject to diverse post-translational modifications (PTMs)^22–26^.

Although previous studies have explored potential links between the DDR and mTORC1 signaling, the mechanistic understanding of mTORC1 regulation in response to genotoxic stress remains incomplete^27^. Here, we report that genotoxic stress alters mTORC1 signaling in a substrate-specific manner, with decreased phosphorylation of TFEB and increased phosphorylation of S6K. In particular, we identify RHEB as a novel ATM substrate whose phosphorylation on specific residues is enhanced by DNA damage. Notably, this modification selectively modulates lysosomal mTORC1 signaling causing reduced TFEB phosphorylation and promoting TFEB activation, thereby coordinating the cellular response to DNA damage. Our findings reveal a regulatory axis coupling DDR with lysosomal mTORC1 signaling through direct modulation of RHEB, and uncover a molecular mechanism that ensures proper adaptation of cells to genotoxic stress via the regulation of growth- and lysosome-related signaling pathways.

## Results

### RHEB is phosphorylated in response to DNA damage

RHEB is best known as a direct activator of mTORC1, with its activity primarily regulated through its nucleotide-binding state under the control of the TSC complex^14,21,26,28^. Despite extensive efforts to identify protein PTMs in an unbiased manner, large-scale phospho-proteomics studies (summarized in the PhosphoSitePlus database; www.phosphosite.org)^29^ have identified only a limited number of phosphorylated residues on human RHEB (Fig. 1A). Among these, the serine at position 175 (S175) was consistently detected as a phosphorylated site across multiple cell lines. Interestingly, many of these phospho-proteomics experiments were specifically designed to search for novel ATM/ATR substrates^29^, suggesting that RHEB may actually be a target of ATM and/or ATR. Given the central role of these kinases in the DNA damage response (DDR), we hypothesized that RHEB^S175^ may be phosphorylated in response to genotoxic stress. To test this, we generated a phospho-specific antibody against RHEB^S175^ and confirmed its specificity using a non-phosphorylatable alanine mutant (Fig. S1A).

**Figure 1.**
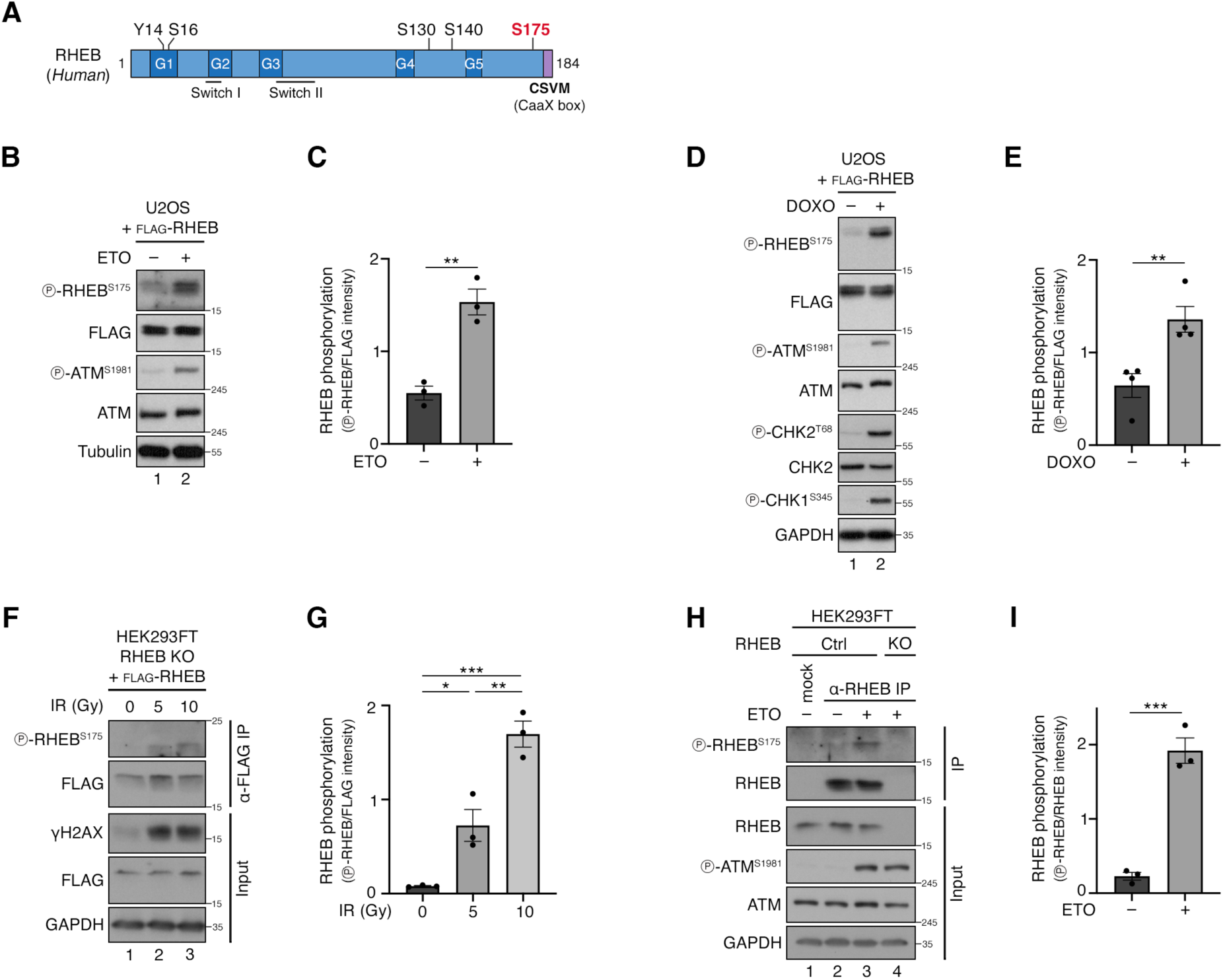
RHEB is phosphorylated at S175 in response to DNA damage. **(A)** Schematic representation of residues on human RHEB identified as phosphorylation sites in phospho-proteomics datasets listed in the PhosphoSitePlus online platform. Also shown is the position of key RHEB elements like the G-boxes 1-5, the Switch I and II regions, and the C-terminal CaaX-box motif. Serine 175 (S175), which is the focus of this study, is highlighted in red. **(B-C)** Immunoblots with lysates from U2OS cells transiently expressing FLAG-tagged RHEB, treated with etoposide (ETO; 20 μM, 2 hr) or DMSO as control, probed with the indicated antibodies. ATM autophosphorylation (ATM^S1981^) was used as a control for DDR induction (B). Quantification of RHEB phosphorylation (p-RHEB/FLAG signal) in (C). n = 3 independent experiments. **(D-E)** Immunoblots with lysates from U2OS cells transiently expressing FLAG-tagged RHEB treated with doxorubicin (DOXO; 2 μM, 2 hr) or DMSO as control, probed with the indicated antibodies. Phosphorylation of ATM (ATM^S1981^), CHK2 (CHK2^T68^) and CHK1 (CHK1^S345^) were used to confirm DDR induction (D). Quantification of RHEB phosphorylation (p-RHEB/FLAG signal) in (E). n = 4 independent experiments. **(F-G)** Immunoblots with lysates from RHEB KO HEK293FT cells stably expressing FLAG-tagged RHEB, left untreated or exposed to increasing doses of ionizing radiation (0-10 Gy), probed with the indicated antibodies (F). Quantification of RHEB phosphorylation (p-RHEB/FLAG signal) in (G). n = 3 independent experiments. **(H-I)** Endogenous RHEB immunoprecipitation from control HEK293FT cells (Ctrl) treated with etoposide (ETO; 20 μM, 2 hr) or DMSO as control, followed by immunoblotting with the indicated antibodies. RHEB KO cells used as negative control (H). Quantification of endogenous RHEB phosphorylation (p-RHEB/RHEB) (I). n = 3 independent experiments. Data in graphs shown as mean ± SEM. * p < 0.05, ** p < 0.01, *** p < 0.001. See also Figure S1.

To determine whether RHEB^S175^ is phosphorylated upon DNA damage, we exposed human osteosarcoma U2OS cells expressing FLAG-tagged RHEB to etoposide, a topoisomerase II inhibitor^30^. As expected, etoposide activated ATM, as indicated by the induction of its autophosphorylation (Fig. 1B)^31^. While basal RHEB phosphorylation was low but detectable, it was markedly increased upon etoposide treatment (Fig. 1B,C). Similar results were obtained following treatment with doxorubicin, another topoisomerase II inhibitor and DNA intercalator (Fig. 1D,E), and with hydroxyurea, which induces replication stress by inhibiting ribonucleotide reductase and thereby depleting deoxyribonucleotide pools (Fig. S1B,C)^32–34^. To assess whether this response extends to other types of genotoxic stress, we irradiated RHEB knock-out (KO) human embryonic kidney HEK293FT cells reconstituted with FLAG-tagged RHEB at 0, 5 or 10 gray (Gy), and observed a dose-dependent increase in RHEB^S175^ phosphorylation (Fig. 1F,G). Similar results were obtained assessing the phosphorylation of endogenous RHEB, immunopurified from HEK293FT cells treated with etoposide (Fig. 1H,I). Together, these results demonstrate that RHEB is phosphorylated at S175 in response to DNA damage induced by diverse stimuli, suggesting a potential regulatory role for this modification in the cellular response to genotoxic stress.

### ATM mediates RHEB phosphorylation in response to DNA damage

Having established that RHEB is phosphorylated upon DNA damage, we next sought to identify the upstream kinase responsible for this modification. In confirmation of the phospho-proteomic datasets that suggested ATM and/or ATR as candidate kinases, we first tested whether RHEB phosphorylation depends on ATM activity. Treatment of cells with the selective ATM inhibitor KU-60019 completely abolished etoposide-induced RHEB S175 phosphorylation (Fig. 2A), implicating ATM as a key regulator. In contrast, inhibition of ATR or DNA-PK did not prevent the increase in RHEB phosphorylation induced by etoposide (Fig. 2B). Consistent with the pharmacological inhibition data, genetic knockdown of ATM, but not ATR, suppressed the etoposide-induced RHEB phosphorylation (Fig. 2C). Together, these results indicate that RHEB phosphorylation in response to DNA damage is predominantly mediated by ATM.

**Figure 2.**
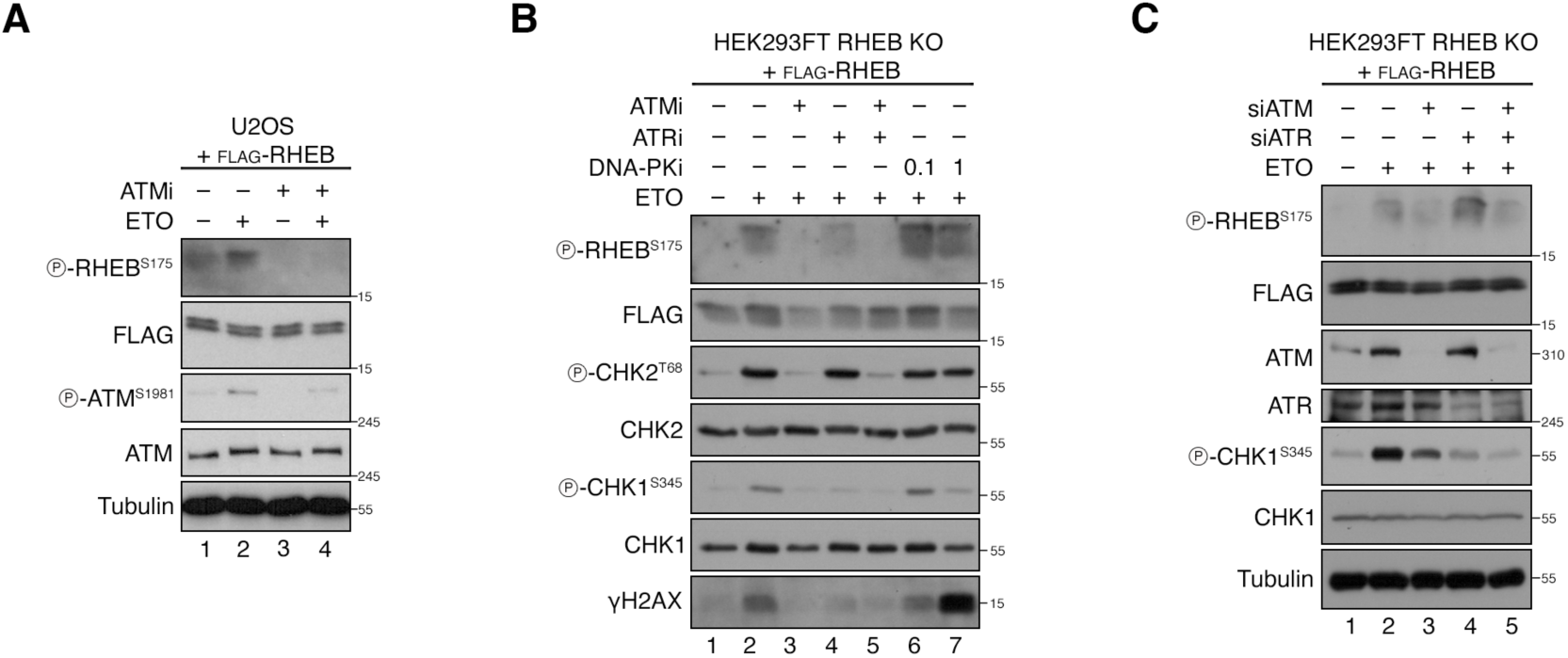
RHEB phosphorylation in response to DNA damage is mediated by ATM. **(A)** Immunoblots with lysates from U2OS cells transiently expressing FLAG-tagged RHEB, pre-treated with ATM inhibitor (ATMi; KU-60019, 1 μM, 30 min), followed by treatment with etoposide (ETO; 20 μM, 2 hr), probed with the indicated antibodies. DMSO was used as control for all treatments. n = 3 independent experiments. **(B)** Immunoblots with lysates from RHEB KO HEK293FT cells stably expressing FLAG-tagged RHEB, pre-treated with ATM inhibitor (ATMi; KU-60019; 1 μM), ATR inhibitor (ATRi; VE-832; 1 μM), or DNA-PK inhibitor (DNA-PKi; NU7441; 0.1 μM or 1 μM) for 30 min, followed by treatment with etoposide (ETO; 20 μM, 2 hr), probed with the indicated antibodies. DMSO was used as control for all treatments. n = 3 independent experiments. **(C)** Immunoblots with lysates from RHEB KO HEK293FT cells stably expressing FLAG-tagged RHEB, transfected with siRNAs targeting ATM (siATM), ATR (siATR), or Luciferase as a control, probed with the indicated antibodies. Cells were treated with etoposide (ETO; 20 μM, 2 hr), or DMSO as control, prior to lysis.

### ATM phosphorylates RHEB directly and independently of the TSC

ATM was previously shown to inhibit mTORC1 activity via the LKB1-AMPK-TSC2 axis in response to oxidative stress^35,36^. Given that the TSC complex, whose core component, TSC2, is the cognate GAP (GTPase-activating protein) for RHEB, we asked whether ATM regulates RHEB phosphorylation indirectly, acting through TSC. To address this, we examined RHEB phosphorylation upon TSC2 knockdown or in TSC1 KO cells following etoposide treatment. In both settings, RHEB phosphorylation was robustly induced by etoposide, indicating that this modification occurs downstream of the TSC complex and independently of the previously-reported ATM-LKB1-AMPK-TSC2 signaling axis (Fig. 3A,B). Together with the observation that RHEB^S175^ constitutes a putative ATM-regulated phospho-site, these results also suggested that ATM may be phosphorylating RHEB directly. Indeed, co-immunoprecipitation (co-IP) experiments revealed the interaction between exogenously expressed FLAG-tagged RHEB and endogenous ATM protein (Fig. 3C), in line with a previous report^26^ suggesting interaction between RHEB and a catalytic domain of ATM. Finally, *in vitro* kinase (IVK) assays using immunopurified FLAG-tagged ATM protein and recombinant RHEB confirmed that ATM is able to phosphorylate RHEB directly and specifically, as the phosphorylation was abolished in the absence of ATP, in the presence of an ATM inhibitor, or using a kinase-dead ATM mutant (Fig. 3D).

**Figure 3.**
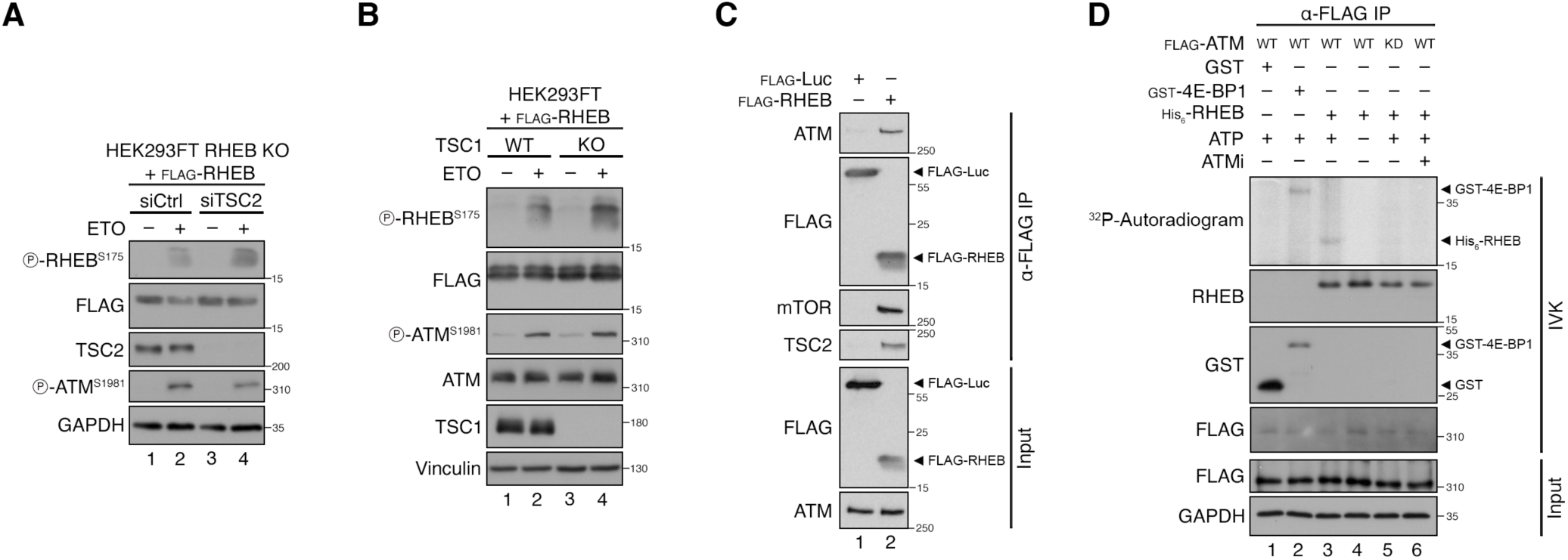
ATM phosphorylates RHEB *in vitro* and independently of TSC. **(A)** Immunoblots with lysates from RHEB KO HEK293FT cells stably expressing FLAG-tagged RHEB, transfected with siRNAs targeting TSC2 (siTSC2) or Luciferase as a control (siCtrl), probed with the indicated antibodies. Cells were treated with etoposide (ETO; 20 μM, 2 hr), or DMSO as control, prior to lysis. **(B)** Immunoblots with lysates from HEK293FT WT or TSC1 KO cells transiently expressing FLAG-tagged RHEB, treated with etoposide (ETO; 20 μM, 2 hr) or DMSO as control, probed with the indicated antibodies. **(C)** Co-immunoprecipitation between endogenous ATM and RHEB from HEK293FT cells transiently expressing FLAG-tagged RHEB or Luciferase (Luc) as a control. The input and IP samples were analyzed by immunoblotting with the indicated antibodies. **(D)** *In vitro* kinase assays with ATM immunopurified from HEK293FT cells transiently expressing FLAG-tagged WT or kinase-dead (KD) ATM, using recombinant His_6_-tagged RHEB protein as a substrate. Cells were treated with etoposide (ETO; 20 μM, 2 hr) before lysis to activate ATM. Specificity of RHEB phosphorylation was confirmed by pre-treatment with an ATM inhibitor (ATMi; KU-60019; 1 μM, 30 min), or omitting ATP from the reaction. Recombinant GST or GST-4E-BP1 proteins were used as a negative or positive control, respectively. Substrate phosphorylation detected by autoradiography. n = 3 independent experiments.

### Substrate-specific mTORC1 responses to DNA damage and RHEB phosphorylation

mTORC1 regulates a broad array of downstream targets to coordinate cellular growth and stress responses. Among these are S6K, which is involved in the regulation of protein synthesis, and TFEB, a transcription factor that governs lysosome biogenesis and autophagy^16–18,20^. We and others have recently shown that mTORC1 activity is not uniformly applied to all substrates, but instead displays substrate-specific regulation. For instance, TFEB—a lysosomal mTORC1 substrate—is regulated distinctly from non-lysosomal substrates such as S6K, especially upon perturbation of lysosomal function or genetic inactivation of the Rag GTPases^17^. On the other hand, hyperactivation of mTORC1 following TSC loss induces S6K hyperphosphorylation while strongly reducing TFEB phosphorylation, thereby promoting TFEB activation and lysosomal biogenesis^37–39^. Accordingly, RHEB KO cells exhibit a reversed pattern, that is diminished S6K phosphorylation and increased TFEB phosphorylation^40^.

To investigate how mTORC1 responds to DNA damage in our setting, we examined the phosphorylation of different substrates following etoposide treatment. In control cells, DNA damage led to a marked reduction in TFEB phosphorylation, while S6K phosphorylation was consistently increased (Fig. 4A-C). Importantly, the dephosphorylation of TFEB was strongly attenuated in RHEB KO cells, indicating that the TFEB response to DNA damage is dependent on RHEB (Fig. 4A-C). S6K phosphorylation, on the other hand, was already minimal in RHEB KO cells and remained low regardless of genotoxic stress. Since the phosphorylation status of TFEB governs its subcellular localization—dephosphorylated TFEB relocalizes to the nucleus to activate target genes—we next assessed TFEB localization following etoposide treatment. In line with its phosphorylation profile, the nuclear localization of TFEB was strongly enhanced in wild-type cells upon DNA damage, whereas this response was diminished in RHEB KO cells (Fig. 4D,E). To determine whether this nuclear translocation correlates with TFEB-mediated lysosomal gene activation, we measured lysosomal abundance using LysoTracker staining. Etoposide treatment led to a robust increase in lysosomal signal in wild-type cells, which was absent in RHEB KO cells (Fig. 4F,G). These findings suggest that TFEB activation following DNA damage promotes lysosomal biogenesis and that this adaptive response is impaired in the absence of RHEB. Together, our data reveal that genotoxic stress triggers TFEB activation and lysosomal biogenesis in a RHEB-dependent manner, while S6K phosphorylation is regulated independently under these conditions.

**Figure 4.**
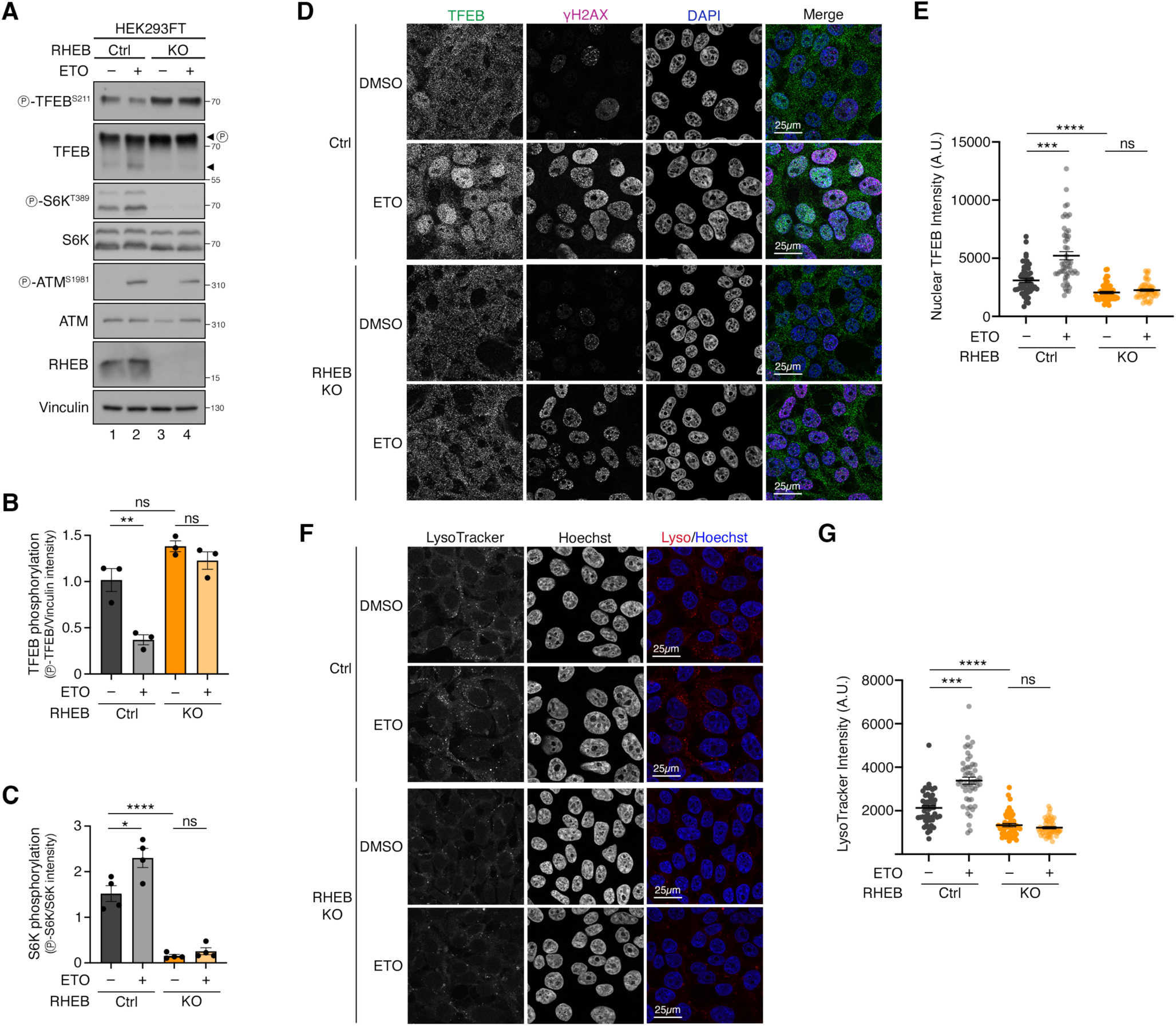
DNA damage induces TFEB activation in a RHEB-dependent manner. **(A-C)** Immunoblots with lysates from control (Ctrl) and RHEB KO HEK293FT cells treated with etoposide (ETO; 20 μM, 8 h) or DMSO as control, probed with the indicated antibodies. Arrowheads indicate bands corresponding to phosphorylated (P) and non-phosphorylated TFEB forms (A). Quantification of TFEB phosphorylation (p-TFEB/loading control) in (B) and S6K phosphorylation (p-S6K/total S6K) in (C). n = 4 independent experiments. **(D-E)** TFEB localization analysis in control (Ctrl) and RHEB KO HEK293FT cells treated as in (A), using confocal microscopy. γ-H2AX staining used as a control for DDR induction. Nuclei stained with DAPI (D). Quantification of nuclear TFEB signal intensity (arbitrary units, A.U.) in (E). n = 50 individual cells from 5 independent fields per condition. **(F-G)** LysoTracker staining in control (Ctrl) and RHEB KO HEK293FT cells treated as in (A), using confocal microscopy. Nuclei stained with Hoechst (F). Quantification of LysoTracker signal intensity (arbitrary units, A.U.) in (G). n = 50 cells from 4-5 independent fields per condition. Scale bars = 25 μm. Data in graphs shown as mean ± SEM. * p < 0.05, *** p < 0.001, **** p < 0.0001; ns, non-significant.

### Phosphorylation of RHEB by ATM is required for TFEB activation following DNA damage

To further investigate the role of RHEB phosphorylation in the DNA damage response, we extended our kinase-substrate prediction analyses beyond publicly available experimental phospho-proteomics data (PhosphoSitePlus) by querying NetworKIN, an online tool that integrates sequence motifs and protein-protein interaction networks to infer kinase-substrate relationships^41,42^. In addition to RHEB S175, NetworKIN predicted serine 6 (S6) as an additional putative ATM-regulated phosphorylation site on RHEB (Table S1). To assess the functional relevance of these sites, we generated a double phospho-deficient RHEB mutant (S6A/S175A; hereafter, ‘SSAA’) and reconstituted RHEB KO HEK293FT cells with either wild-type (RHEB^WT^) or SSAA mutant RHEB (RHEB^SSAA^). Stable re-expression of either RHEB^WT^ or RHEB^SSAA^ restored basal mTORC1 signaling in RHEB KO cells, as evidenced by recovery of S6K phosphorylation (Fig. S2), indicating that the RHEB^SSAA^ mutant retains activity under steady-state conditions.

We next examined how these cells respond to DNA damage. Upon etoposide treatment, RHEB^WT^ cells showed a clear decrease in TFEB phosphorylation, whereas this response was strongly attenuated in RHEB^SSAA^-expressing cells (Fig. 5A,B). Notably, DNA damage induction similarly increased S6K phosphorylation in both RHEB^WT^ and RHEB^SSAA^ cells, suggesting that the defect is specific to the regulation of lysosomal mTORC1 signaling and TFEB phosphorylation (Fig. 5A,C). Consistent with the signaling data, TFEB nuclear translocation was induced by DNA damage in RHEB^WT^ cells but this response was impaired in RHEB^SSAA^-expressing cells (Fig. 5D,E). Similarly, etoposide-induced lysosomal biogenesis—monitored by LysoTracker staining—was evident in RHEB^WT^-reconstituted cells but absent in RHEB^SSAA^-expressing cells (Fig. 5F,G). Together with our earlier results in RHEB KO cells, these data support a model based on which ATM-dependent phosphorylation of RHEB is essential for engaging the TFEB axis downstream of mTORC1 during DDR activation.

**Figure 5.**
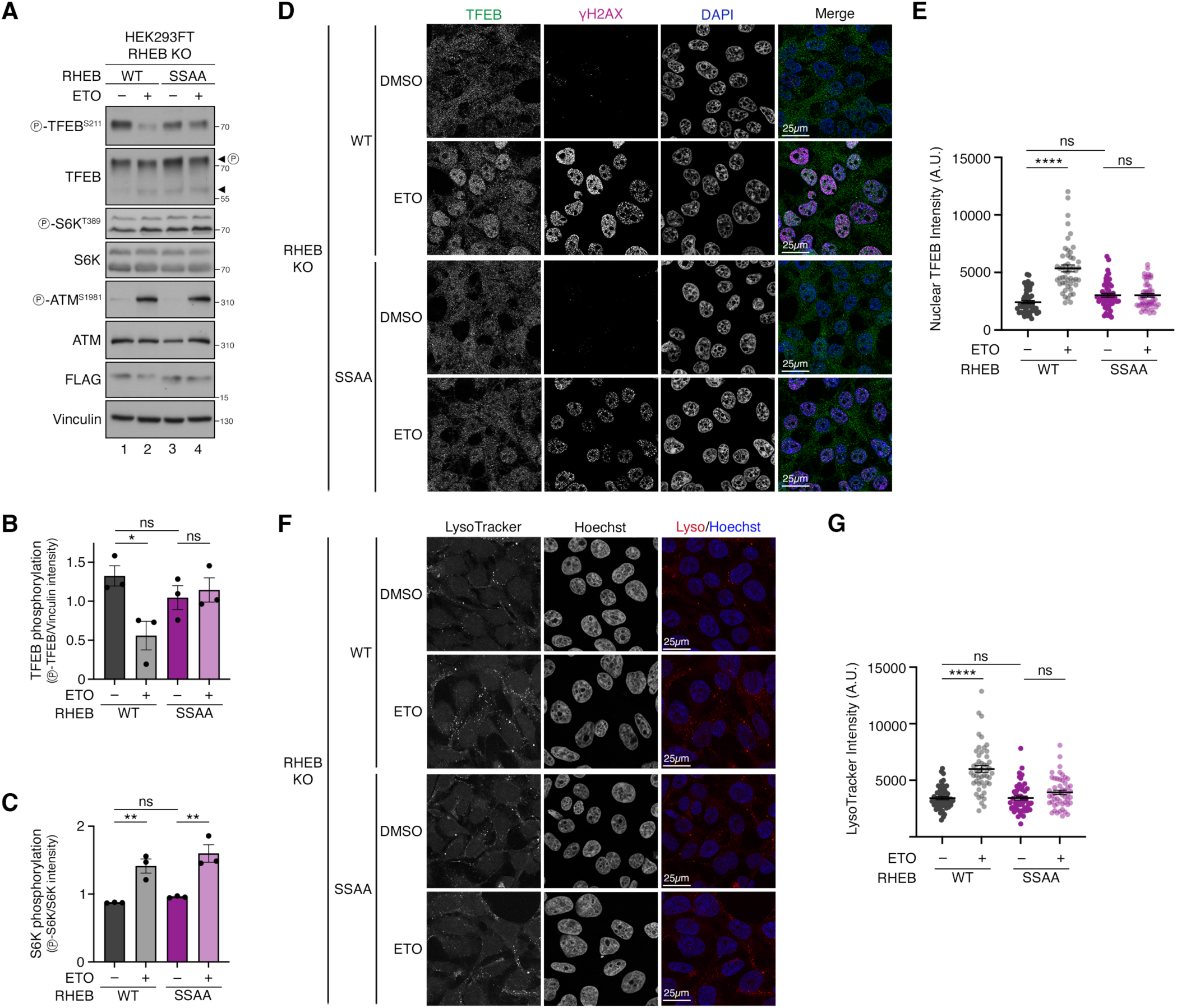
RHEB phosphorylation by ATM is required for TFEB activation in response to DNA damage. **(A-C)** Immunoblots with lysates from RHEB KO HEK293FT cells stably expressing FLAG-tagged wild-type RHEB (WT) or a phosphorylation-deficient RHEB^S6A/S175A^ mutant (SSAA), treated with etoposide (ETO; 20 μM, 8 h) or DMSO as control, probed with the indicated antibodies. Arrowheads indicate bands corresponding to phosphorylated (P) and non-phosphorylated TFEB forms (A). Quantification of TFEB phosphorylation (p-TFEB/loading control) in (B) and S6K phosphorylation (p-S6K/total S6K) in (C). n = 3 independent experiments. **(D-E)** TFEB localization analysis in RHEB KO HEK293FT cells stably expressing FLAG-tagged wild-type RHEB (WT) or a phosphorylation-deficient RHEB^S6A/S175A^ mutant (SSAA) treated as in (A), using confocal microscopy. γ-H2AX staining used as a control for DDR induction. Nuclei stained with DAPI (D). Quantification of nuclear TFEB signal intensity (arbitrary units, A.U.) in (E). n = 50 cells from 4-5 independent fields per condition. **(F-G)** LysoTracker staining in RHEB KO HEK293FT cells stably expressing FLAG-tagged RHEB WT (WT) and FLAG-tagged RHEB^S6A/S175A^ (SSAA) treated as in (A) using confocal microscopy. Nuclei stained with Hoechst (F). Quantification of LysoTracker signal intensity (arbitrary units, A.U.) in (G). n = 50 cells from 4-5 independent fields per condition. Scale bars = 25 μm. Data in graphs shown as mean ± SEM. ** p < 0.01, **** p < 0.0001; ns, non-significant. See also Figure S2.

### TFEB activation facilitates recovery of cell growth following DNA damage

Given the role of mTORC1 in adapting cellular physiology to environmental cues, we asked whether RHEB-mediated TFEB activation contributes to the cellular response to DNA damage. To assess this, we monitored cell growth dynamics following ionizing radiation by measuring confluency over time. Wild-type cells exhibited a transient pause in proliferation after irradiation (10 Gy), followed by recovery and continued growth (Fig. 6A). In contrast, RHEB KO cells showed a clear growth defect under the same conditions, following the initial expansion phase. This effect was specific to the cellular response to irradiation, as RHEB KO cells displayed similar growth kinetics to wild-type controls under basal conditions (Fig. S3A). Similarly, cells expressing the non-phosphorylatable RHEB^SSAA^ mutant were more sensitive to irradiation than RHEB^WT^ cells (Fig. 6C), despite comparable growth between the two genotypes under unstressed conditions (Fig. S3B), again showing that RHEB phosphorylation is a key event that is required for proper cellular response to genotoxic stress.

**Figure 6.**
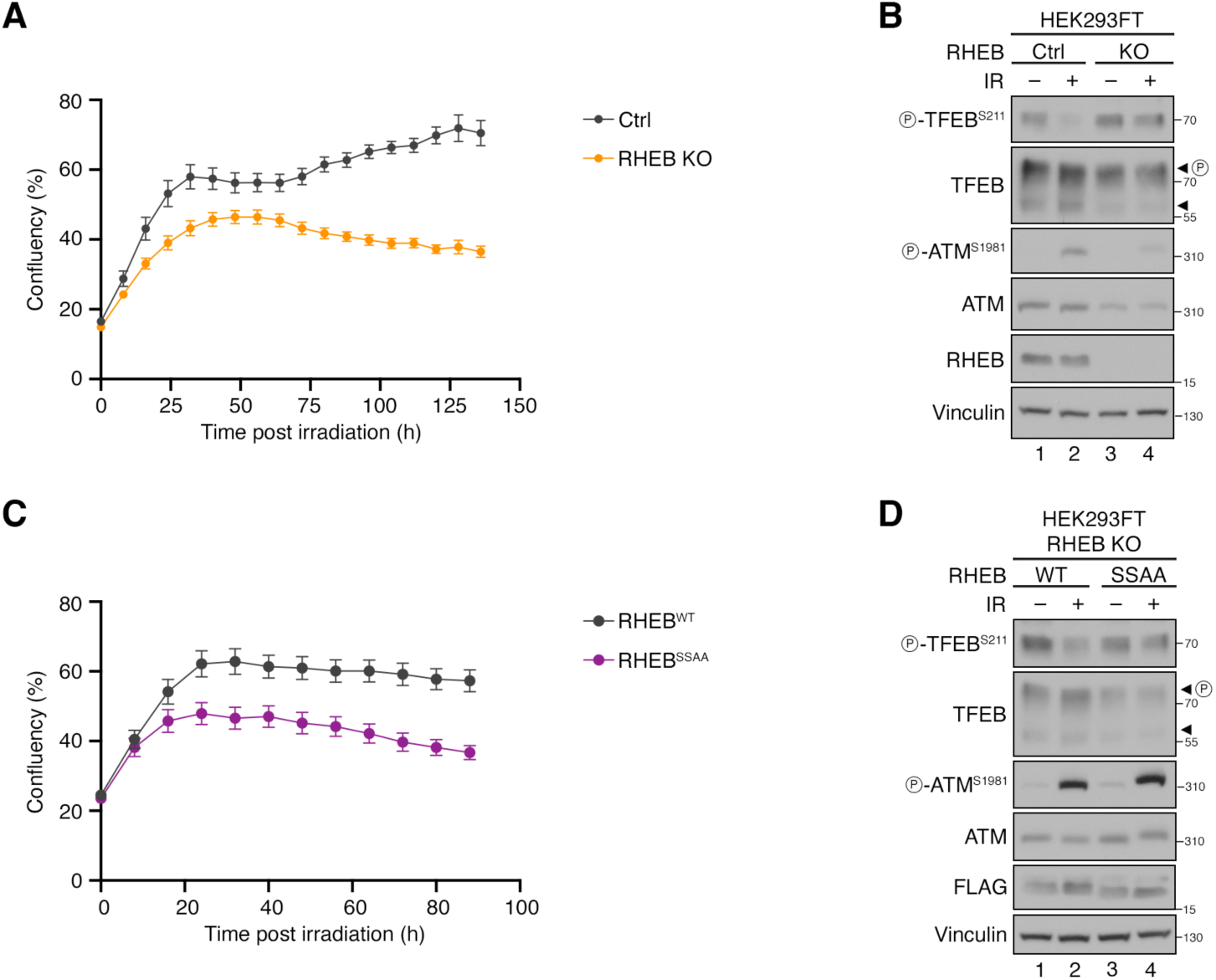
RHEB phosphorylation is required for recovery of cell growth following DNA damage. **(A)** Time-course of cell confluency (%) with control (Ctrl) and RHEB KO HEK293FT cells, following ionizing radiation (IR; 10 Gy). **(B)** Immunoblots with lysates from control (Ctrl) and RHEB KO cells irradiated with 10 Gy for 8 h (IR) or left untreated as control. Decrease in TFEB dephosphorylation is observed in control cells following irradiation, but not in RHEB KOs. **(C)** Time-course of cell confluency (%) with RHEB KO HEK293FT cells stably expressing FLAG-tagged wild-type RHEB (WT) or a phosphorylation-deficient RHEB^S6A/S175A^ mutant (SSAA), following irradiation (IR; 20 Gy). **(D)** Immunoblots with lysates from RHEB KO HEK293FT cells stably expressing FLAG-tagged wild-type RHEB (WT) or a phosphorylation-deficient RHEB^S6A/S175A^ mutant (SSAA), irradiated with 20 Gy (IR) for 8 h or left untreated as control. Decrease in TFEB phosphorylation is observed in RHEB^WT^-expressing cells, a response that is blunted in RHEB^SSAA^-expressing cells. See also Figures S3-S4.

Since both RHEB KO and RHEB^SSAA^ cells exhibit impaired TFEB activation upon DNA damage (Fig. 6B,D), we speculated that the deficient TFEB response may underlie their impaired growth recovery. In contrast, our data indicate that this effect is unlikely to involve S6K, as its enhanced phosphorylation upon etoposide treatment remained unaffected in RHEB^SSAA^-reconstituted cells (Fig. 5A-C). In support of our hypothesis, prior studies have implicated TFEB in mediating DDR and DNA repair pathways^43–46^. To explore this possibility further, we performed gene ontology (GO) term enrichment analysis using a curated list of TFEB target genes. As expected, the analysis revealed enrichment of canonical TFEB-regulated biological processes such as lysosome organization (GO:0007040), autophagy (GO:0006914), and mTORC1 signaling (GO:1904263) (Fig. S4). Interestingly, we also observed significant enrichment of biological processes linked to the DDR, including DNA repair (GO:000628), DNA damage checkpoint signaling (GO:0000077, GO:0006974, GO:0042770), response to radiation (GO:0071478), and cell cycle regulation (GO:0051726, GO:0000082), among others (Fig. S4).

To directly test whether the impaired TFEB response contributes to the growth phenotype, we manipulated the Rag GTPases, which tether TFEB to lysosomes and restrict its nuclear translocation. Importantly, S6K phosphorylation is not affected by the presence or activation status of the Rags, when exogenous amino acids are present^10,17^. To interrogate if the compromised physiological response of RHEB KO cells to irradiation can be attributed to the blunted TFEB activation, we transiently depleted RagA and RagB to release TFEB from the lysosomes. As expected, knockdown of RagA/B in RHEB KO cells resulted in loss of TFEB dephosphorylation (Fig. 7A). Strikingly, this perturbation also fully rescued the irradiation-driven growth impairment of RHEB KO cells (Fig. 7B). Conversely, expression of a constitutively active RagA/C dimer in wild-type cells, which caused sustained TFEB phosphorylation even in irradiated cells (Fig. 7C) and is known to sequester TFEB on lysosomes^18,20,47^, phenocopied the impaired growth seen in RHEB KO cells following irradiation (Fig. 7D). Again, expression of active RagA/C or RagA/B knockdown only marginally affected cell growth under non-stressed conditions (Fig. S5A,B), reinforcing the role of TFEB regulation in modulating the cellular adaptation to genotoxic stress.

**Figure 7.**
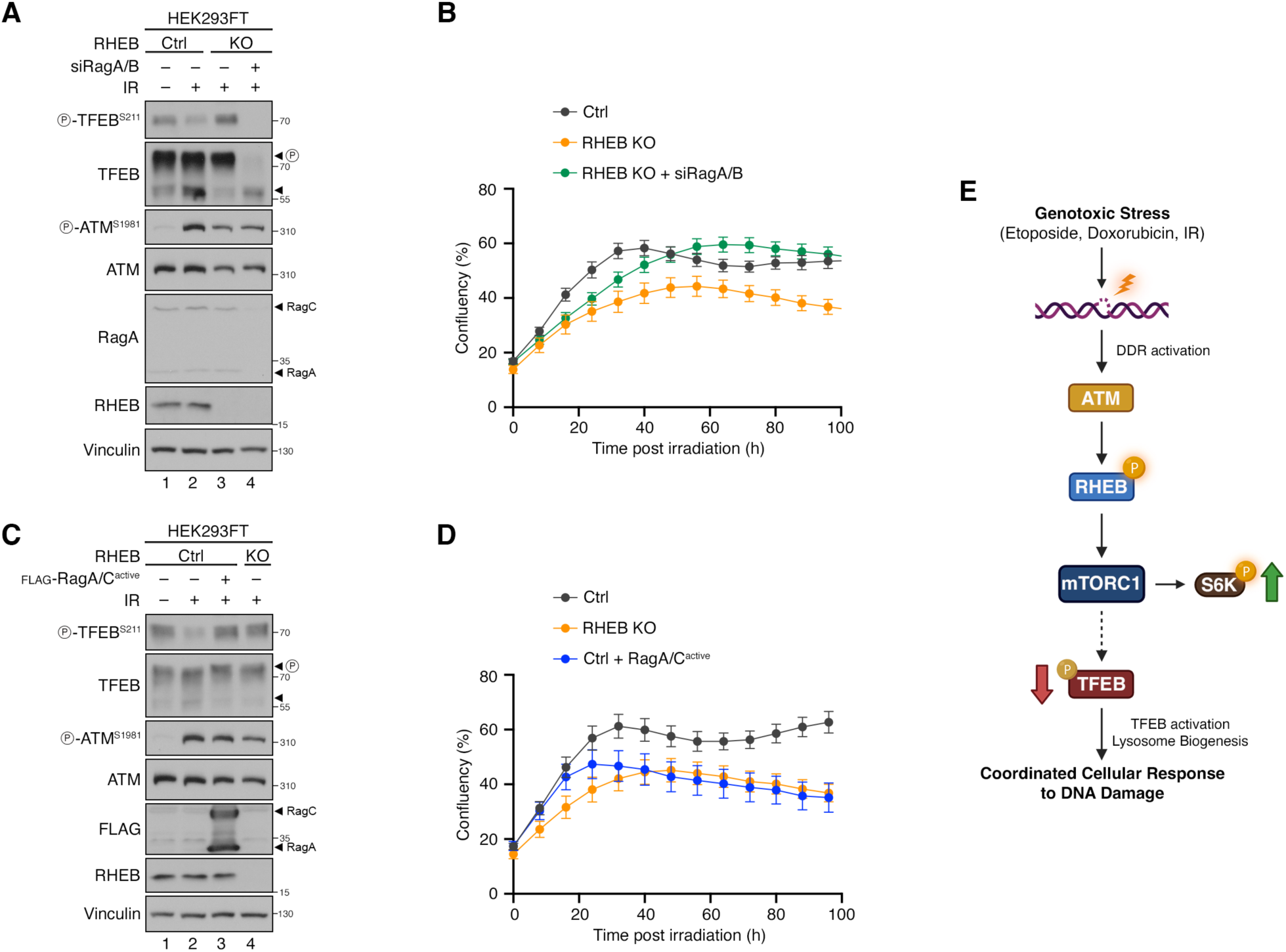
RHEB-phosphorylation-dependent activation of TFEB is required for recovery of cell growth following DNA damage. **(A)** Immunoblots with lysates from controls (Ctrl), RHEB knockouts (RHEB KO), or RHEB KO cells with RagA/B knockdown (RHEB KO + siRagA/B), irradiated with 10 Gy for 8 h (IR) or left untreated as control. RagA/B knockdown abolishes TFEB phosphorylation, as expected. **(B)** Time-course of HEK293FT cell confluency (%) with controls (Ctrl), RHEB knockouts (RHEB KO), or RHEB KO cells with RagA/B knockdown (RHEB KO + siRagA/B), following irradiation (10 Gy). RagA/B knockdown rescues the growth defect upon irradiation in RHEB KO cells. **(C)** Immunoblots with lysates from controls (Ctrl), RHEB knockouts (RHEB KO), or cells transiently expressing active RagA/C dimers (Ctrl + RagA/C^active^), irradiated with 10 Gy for 8 h (IR) or left untreated as control. Expression of active RagA/C dimers prevents TFEB dephosphorylation upon irradiation. **(D)** Time-course of HEK293FT cell confluency (%) with controls (Ctrl), RHEB knockouts (RHEB KO), or cells transiently expressing active RagA/C dimers (Ctrl + RagA/C^active^), following irradiation (10 Gy). **(E)** Schematic model of the ATM-RHEB-mTORC1-TFEB signaling axis activated in response to genotoxic stress. DNA damage induced by etoposide, doxorubicin or ionizing radiation (IR) activates ATM, which directly phosphorylates RHEB. This modification attenuates mTORC1 activity toward TFEB, causing TFEB dephosphorylation and nuclear translocation, while phosphorylation of S6K remains elevated. TFEB activation enhances lysosomal biogenesis in a RHEB phosphorylation-dependent manner. A proper TFEB response is essential for orchestrating the cellular response to genotoxic stress (see also text for details). See also Figure S5.

Collectively, these results establish TFEB activation as a critical component of the post-DDR recovery program, and demonstrate that this process requires the ATM-dependent phosphorylation of RHEB. Hence, our findings support a model in which RHEB-mTORC1-TFEB signaling enables adaptation programs that contribute to cellular recovery from genotoxic insult.

## Discussion

### RHEB integrates multiple stimuli to regulate mTORC1 activity

RHEB is a well-established direct activator of mTORC1, with its activity primarily regulated through its GTP/GDP loading state that is under the control of the TSC complex. Most upstream cues that impinge on mTORC1 activity—including amino acids, growth factors, and energy status—converge on the TSC-RHEB axis. More recently, PTMs such as ubiquitination and phosphorylation have emerged as additional layers of RHEB regulation, enabling finer control over its activity. For example, PRAK (p38-regulated/activated protein kinase; also known as MAPK5)-mediated phosphorylation of RHEB at S130 reduces its activity under energetic stress^25^. In this study, we identify a novel ATM-mediated phosphorylation event that is induced in response to DNA damage. This modification is consistently observed across treatments with DNA-damaging agents that target distinct molecular mechanisms, as well as ionizing radiation (Fig. 1 and Fig. S1). Importantly, this phosphorylation occurs independently of the TSC complex, positioning RHEB as a direct substrate of ATM. These findings reveal a previously unrecognized role for RHEB in fine-tuning mTORC1 activity in response to genotoxic stress, and unfold an alternative mechanism by which DNA damage signals can influence mTORC1 activity.

Previous reports showed that oxidative and nitrosative stresses can inhibit mTORC1 via an ATM-LKB1-AMPK-TSC2 axis^35,36^. Unlike these studies that showed a TSC-dependent modulation of mTORC1 activity downstream of ATM, our data demonstrate that this kinase can also phosphorylate RHEB directly in response to DNA damage (Fig. 3). Together with prior work identifying various PTMs on RHEB—such as ubiquitination, NEDDylation, and phosphorylation—our findings further expand the role of RHEB beyond being an effector of the TSC complex. Thus, RHEB emerges as a dynamic signaling hub that integrates diverse cellular cues to regulate mTORC1 activity^22–25^. By identifying ATM as an upstream kinase that directly modifies RHEB upon genotoxic stress, our study underscores the versatility of RHEB as a regulatory node in the broader mTORC1 signaling network.

### DNA damage is signaled through the ATM-RHEB axis to drive substrate-specific mTORC1 regulation

Our findings reveal a substrate-specific rewiring of mTORC1 signaling in response to genotoxic stress. Upon DNA damage, TFEB phosphorylation is reduced, leading to its nuclear translocation and activation of lysosomal biogenesis programs. In contrast, phosphorylation of canonical, non-lysosomal substrates such as S6K is increased. This decoupling of mTORC1 outputs supports a model in which the pathway is not simply turned ‘on’ or ‘off’, but rewired toward specific substrates depending on the context.

Mechanistically, we report that this selective regulation is dependent on RHEB phosphorylation by ATM. Cells lacking RHEB or expressing a non-phosphorylatable RHEB mutant fail to exhibit TFEB dephosphorylation or nuclear translocation in response to DNA damage. As a result, these cells show a markedly impaired ability to increase lysosomal content upon genotoxic stress, and display delayed growth recovery following irradiation. In contrast, the S6K phosphorylation response to DNA damage remained intact in RHEB^SSAA^-expressing cells, indicating that the defect is specific to the lysosomal signaling branch of mTORC1.

Functionally, the cells with attenuated TFEB response (i.e., RHEB KO, RHEB^SSAA^ mutants, or cells expressing active RagA/C dimers) show compromised recovery in cell confluency following DNA damage. Conversely, cells with functional RHEB-phosphorylation-dependent TFEB activation manage genomic stress more effectively by transiently arresting, presumably initiating repair, and resuming proliferation. This observation builds on the concept of substrate-specific mTORC1 regulation and adds a genotoxic dimension that implicates RHEB phosphorylation as a conditional switch that selectively regulates the lysosomal mTORC1-TFEB axis during DNA damage while sparing cytosolic mTORC1-S6K signaling.

These findings align with the growing evidence that mTORC1 activity is not uniformly regulated across its substrates^10,17,37–40,48–53^. We and others have previously demonstrated that mTORC1 exhibits substrate-specific regulation, particularly evident when comparing lysosomal versus non-lysosomal targets. For instance, phosphorylation of lysosomal mTORC1 substrates, such as TFEB and TFE3, requires intact Rag GTPase dimers and proper lysosomal function, while canonical cytosolic substrates like S6K and 4E-BP1 are largely Rag- and lysosome-independent under basal culture conditions^17^. As a result, genetic or pharmacological perturbation of lysosomal mTORC1 selectively affects the lysosomal branch, sparing non-lysosomal targets^17,52^. Similar patterns are observed in cells expressing cancer-related activating RagC mutants, which markedly enhance TFEB phosphorylation without strongly affecting S6K^10^. Furthermore, in TSC-null models, cytosolic targets like S6K are hyperphosphorylated, while TFEB/TFE3 are hypophosphorylated^37–39^, whereas RHEB KO cells display the reverse phenotype^40^.

Our study extends these observations to DNA damage responses, where mTORC1 exhibits substrate-specific regulation of its lysosomal and cytosolic substrates. Furthermore, ATM-mediated phosphorylation of RHEB is required for TFEB regulation upon DNA damage, while the S6K response remains unaffected in the absence of RHEB phosphorylation. Together, our findings provide evidence for a DNA-damage-responsive, RHEB-phosphorylation-dependent regulatory axis that selectively engages the lysosomal arm of mTORC1 signaling without affecting its canonical, cytosolic targets. This reveals an additional layer of control through which mTORC1 integrates metabolic and genotoxic cues and reinforces the emerging view that mTORC1 functions as a substrate-selective signaling node, rather than a binary switch.

### Fine-tuning of mTORC1 activity through the ATM-RHEB axis is important for proper cellular response to DNA damage

Previous studies investigating the effect of DNA damage on mTORC1 signaling have yielded mixed results, likely due to differences in cell lines, experimental conditions, DNA-damage-inducing protocols, or the read-outs used to assess mTORC1 activity^27,43,45,54–56^. While some reports observed suppression of mTORC1 following DNA damage, others noted maintained or increased activity toward substrates like S6K. Our findings help reconcile these differences by demonstrating that mTORC1 activity is regulated in a substrate-specific manner upon DNA damage. While lysosomal targets like TFEB are dephosphorylated and activated, the phosphorylation of cytosolic targets like S6K is further enhanced. These observations are consistent with prior studies showing increased S6K activity following genotoxic stress and implicating S6K in several DDR-related functions, such as supporting G2/M arrest via CDK1 (cyclin dependent kinase 1) to facilitate homologous recombination, enhancing mismatch repair through MSH6 (mutS homolog 6) phosphorylation, and promoting p53 stabilization^55–57^.

In parallel, our data firmly position TFEB as a crucial mediator of cellular adaptation to DNA damage. Multiple studies have shown that TFEB activation promotes lysosomal biogenesis and autophagy, and plays direct roles in DDR by regulating DNA repair, checkpoint activation, cell cycle arrest, and apoptosis^43–46^. This multifaceted TFEB-dependent response occurs through the regulation of multiple key components in these processes, such as p21, MDM2 and p53^43–46^. The impaired cellular recovery that we observe in models with defective TFEB activation (e.g., in RHEB KO, RHEB^SSAA^, or active RagA/C-expressing cells) further supports this role. Conversely, releasing TFEB from lysosomal retention (e.g., by RagA/B knockdown) restores proliferation even in a RHEB-deficient context. Furthermore, the fact that TFEB activation is accompanied by increased lysosomal abundance suggests a potential role for lysosomal function and cellular recycling processes in supporting DDR. This notion is supported by studies demonstrating that lysosomes and autophagy are critical for DDR protein recruitment to DNA lesions and efficient DNA repair^58–63^.

mTORC1 signaling and the DNA damage response are both tightly linked to ageing and age-related disorders, primarily through their dysregulation. Imbalances in these pathways contribute to cancer, neurodegeneration and other age-associated conditions. Therapeutic targeting of mTORC1 and DDR components is currently being explored in various clinical contexts, particularly in cancer. Within this framework, our study provides additional mechanistic insight by identifying a novel axis through which DNA damage modulates mTORC1 activity: via ATM-mediated phosphorylation of RHEB and subsequent TFEB regulation. This underscores RHEB phosphorylation as a potential marker to study the induction of genotoxic stress in cells. In summary, our study highlights a previously unrecognized mode of mTORC1 regulation by genotoxic stress, adding to the mechanistic understanding of how these two major pathways interact in contexts relevant to ageing and human disease.

## STAR Methods

### EXPERIMENTAL MODEL AND STUDY PARTICIPANT DETAILS

#### Cell culture

All cell lines were maintained at 37 °C in a humidified incubator with 5% CO_2_. Human female embryonic kidney HEK293FT cells (#R70007, Invitrogen; RRID: CVCL_6911) were cultured in high-glucose Dulbecco’s Modified Eagle Medium (DMEM; #41965039, Gibco) supplemented with 10% fetal bovine serum (FBS; #F7524, Sigma; #P30-3306, PAN-Biotech; #FBS.HP.0500, Bio&SELL) and 1x Penicillin-Streptomycin (#15140122, Gibco; #P4333-100ML, Sigma). Human osteosarcoma U2OS cells (#HTB-96, ATCC; RRID: CVCL_0042) were cultured in high-glucose DMEM GlutaMAX (#61965026, Gibco) with 10% FBS and 1x Penicillin-Streptomycin.

HEK293FT cells were purchased from Invitrogen, and U2OS cells were generously provided by Nils-Göran Larsson (MPI-AGE). The identity of the HEK293FT cells was validated using the Multiplex human Cell Line Authentication test (Multiplexion GmbH), which employs a single nucleotide polymorphism (SNP) typing approach, as described at www.multiplexion.de. No commonly misidentified cell lines were used in this study. All cell lines were regularly tested for *Mycoplasma* contamination using a PCR-based method and were confirmed to be *Mycoplasma*-free.

#### Plasmid DNA transfections

Plasmid DNA transfections were performed using Effectene transfection reagent (#301425, QIAGEN), according to the manufacturer’s instructions.

#### Generation of knockout cell lines

The HEK293FT RHEB and TSC1 knockout cell lines were described previously^40,64^.

#### Generation of stable cell lines

Polyclonal RHEB KO HEK293FT cells stably expressing FLAG-tagged RHEB WT or S6A/S175A (SSAA) were generated using a doxycycline-inducible sleeping-beauty-based transposon system^7,65^. RHEB KO cells were co-transfected with pITR-FLAG-RHEB WT or SSAA expression constructs and the pCMV-Trp transposase vector (10:1 ratio). Thirty-six hours post-transfection, cells were selected with 2 μg/ml puromycin (#A1113803, Gibco). To test successful integration of transposons into the genome of the resulting cell lines, expression was induced with varying concentrations of doxycycline (#D9891, Sigma) and monitored by immunoblotting. All experiments were performed without doxycycline induction, as leaky expression levels were comparable across constructs and to endogenous RHEB level.

#### Gene silencing experiments

Transient knockdown of *ATM, ATR, TSC2, RRAGA* and *RRAGB* was performed using siGENOME (pool of 4) gene-specific siRNAs (Horizon Discoveries). An siRNA duplex targeting the *R. reniformis* luciferase gene (RLuc) (#P-002070-01-50, Horizon Discoveries) was used as a control. Transfections were performed using 20 nM siRNA and the Lipofectamine RNAiMAX transfection reagent (#13778075, Invitrogen), according to the manufacturer’s instructions. Cells were harvested 72 hours post-transfection and knockdown efficiency was verified by immunoblotting.

### METHOD DETAILS

#### Cell culture treatments

For drug treatments, the following concentrations and time points were used unless otherwise specified in figure legends: etoposide (#E1383, Sigma) at 20 µM for 2 or 8 h; doxorubicin (#D1515, Sigma) was applied at 2 µM for 2 h; hydroxyurea (#H8627, Sigma) at 2 mM for 2h. For kinase activity inhibition, cells were pre-treated with KU-60019 (ATM inhibitor; #S1570, Selleckchem; 1 μM) VE-821 (ATR inhibitor; #S8007, Selleckchem; 1 μM), and NU7441 (DNA-PK inhibitor; #S2638, Selleckchem; 0.1 or 1 μM) for 30 minutes prior to addition of DNA-damaging agents. DMSO (#4720.1, Roth) was used as vehicle control for all treatments except for hydroxyurea, for which water was used.

#### Antibodies

The custom-made, rabbit polyclonal phospho-specific antibody recognizing RHEB when phosphorylated at S175 (phospho-RHEB^S175^) was produced by immunizing animals with a synthetic KLH (Keyhole Limpet Hemocyanin)-conjugated phospho-peptide corresponding to residues around S175 of human RHEB: MDGAA(pS)QGKSSC. Peptide synthesis was performed by Peptide Specialty Laboratories GmbH (Heidelberg, Germany), and antibody generation was outsourced to ProSci incorporated (Poway, CA, USA). Antibodies were purified by coupling to NHS-Activated Sepharose 4 Fast Flow resin (#17090601, Cytiva), alternating acid-base washing steps, and elution with 0.2 M glycine (pH 2.8). Eluted fractions were normalized by 0.4 M Na_2_HPO_4_ (pH 8.2), and antibodies were stored in 50% glycerol, 0.1% BSA at −20 °C.

A complete list of all primary and secondary antibodies used in this study can be found in Table S2.

#### Identification of phosphorylation sites on RHEB

Phosphorylation sites on RHEB were identified using curated phospho-proteomics data and computational predictions. A putative phosphorylation site (RHEB^S175^) was retrieved from the PhosphoSitePlus database (www.phosphosite.org)^29^, based on unpublished high-throughput phospho-proteomics data generated by Cell Signaling Technology (CST). Kinase-substrate predictions were performed using NetworKIN^42^, which integrates consensus phosphorylation motifs with contextual information, including protein-protein interactions, co-expression data, and subcellular localization, to predict upstream kinases for candidate phosphorylation sites. Human RHEB (UniProt ID: Q15382) was used as input, and high-confidence predictions were prioritized for further investigation.

#### Plasmids and Molecular Cloning

The pcDNA3-FLAG-RHEB WT and pcDNA3-FLAG-Luc (FLAG-tagged firefly Luciferase; used as a negative control) plasmids were described previously^66^. The pcDNA3-FLAG-RHEB S175A and the pcDNA3-FLAG-RHEB S6A/S175A were generated by site-directed mutagenesis using appropriate oligos, and the resulting PCR products were cloned into the EcoRI and NotI restriction sites of the pcDNA3-FLAG-RHEB WT vector. The FLAG-RHEB WT and FLAG-RHEB S6A/S175A mutant were subcloned into the sleeping-beauty-based, doxycycline-inducible pITR-TTP2 vector (based on pITR-TTP^7,40,65^, with a more expanded multiple cloning site feature) using the PvuII/NotI restriction sites, while also removing the additional start codon from the RHEB cDNA. The pETM11-RHEB used for recombinant His_6_-tagged RHEB protein production was generated by PCR-amplifying human RHEB cDNA using appropriate primers and subcloning into the NcoI/NotI restriction sites of the pETM11 vector.

The pcDNA3.1(+) Flag-His-ATM wt (plasmid #31985) and the pcDNA3.1(+) Flag-His-ATM kd (plasmid #31986) were purchased from Addgene (both deposited by Michael Kastan and described in^67^). The pcDNA3-FLAG-RagA Q66L (GTP-locked) and the pcDNA3-FLAG-RagC S75N (GDP-locked) were described previously^68^. All restriction enzymes were purchased from Fermentas/Thermo Scientific. The integrity of all constructs was verified by sequencing. All DNA oligonucleotides used in this study are listed in Table S3.

#### Cell lysis and immunoblotting

For standard SDS-PAGE and immunoblotting experiments, cells from one well of a 6-well plate were lysed in 300 μl ice-cold Triton lysis buffer (50 mM Tris pH 7.5, 1% Triton X-100, 150 mM NaCl, 50 mM NaF, 2 mM Na-vanadate, 0.011 gr/ml beta-glycerophosphate) or RIPA buffer (50 mM Tris pH 7.5, 1% Triton X-100, 150 mM NaCl, 0.1% SDS, 0.5 % sodium deoxycholate, 50 mM NaF, 2 mM Na-vanadate, 0.011 gr/ml beta-glycerophosphate), supplemented with 1x PhosSTOP phosphatase inhibitors (#04906837001, Roche) and 1x cOmplete protease inhibitors (#11697498001, Roche). Lysates were incubated for 10 minutes on ice and clarified by centrifugation (15000 rpm, 15 min, 4 °C). Protein concentration was measured using Protein Assay Dye Reagent (#5000006, Bio-Rad). Normalized samples were boiled in 1x SDS sample buffer for 5 min at 95 °C (6x SDS sample buffer: 350 mM Tris-HCl pH 6.8, 30% glycerol, 600 mM DTT, 12.8% SDS, 0.12% bromophenol blue).

Protein samples were subjected to electrophoretic separation on SDS-PAGE and analyzed by standard Western blotting techniques. In brief, proteins were transferred to nitrocellulose membranes (#10600002 or #10600001, Amersham) and stained with 0.2% Ponceau solution (#33427-01, Serva) to confirm equal loading. Membranes were blocked with 5% skim milk powder (#42590, Serva) in PBS-T [1x PBS, 0.1% Tween-20 (#A1389, AppliChem)] for 1 hour at room temperature, washed three times for 5 min with PBS-T and then incubated with primary antibodies [in PBS-T, 5% bovine serum albumin (BSA; #10735086001, Roche; #8076, Carl Roth)] overnight at 4 °C. The next day, membranes were washed three times for 5 min with PBS-T and incubated with the appropriate HRP-conjugated secondary antibodies (1:10000 in 5% milk in PBS-T) for 1 hour at room temperature. Signals were detected by enhanced chemiluminescence (ECL), using ECL Western Blotting Substrate (#W1015, Promega); or SuperSignal West Femto Substrate (#34095, Thermo Scientific) for weaker signals. Immunoblot images were captured on films (#28906835, GE Healthcare; #4741019289, Fujifilm). Blots were scanned and then quantified using GelAnalyzer 19.1. A list of all primary and secondary antibodies used in this study is provided in Table S2.

#### Immunoprecipitation and co-immunoprecipitation (co-IP)

For ATM and FLAG-RHEB co-immunoprecipitation experiments, cells from a near-confluent 6-well plate were lysed in 500 μl CHAPS IP buffer (50 mM Tris pH 7.5, 0.3% CHAPS, 150 mM NaCl, 50 mM NaF, 2 mM Na-vanadate, 0.011 gr/ml beta-glycerophosphate) supplemented with 1x PhosSTOP phosphatase inhibitors (#04906837001, Roche) and 1x cOmplete protease inhibitors (#11697498001, Roche) for 10 minutes on ice. For anti-RHEB immunoprecipitation experiments, cells from a near-confluent 10-cm dish were lysed in 1 ml Triton IP buffer (50 mM Tris pH 7.5, 1% Triton X-100, 150 mM NaCl, 50 mM NaF, 2 mM Na-vanadate, 0.011 gr/ml beta-glycerophosphate), supplemented with 1x PhosSTOP phosphatase inhibitors (#04906837001, Roche) and 1x cOmplete protease inhibitors (#11697498001, Roche). Samples were clarified by centrifugation (15000 rpm, 15 min, 4 °C) and a fraction of the samples was taken as input. For anti-FLAG IPs, the remaining supernatants were incubated with 30 μl of pre-washed anti-FLAG M2 affinity gel (#A2220, Sigma) at 4 °C on a rotating mixer for 3 h. For anti-RHEB IPs, the supernatant was incubated with 3 µl anti-RHEB antibody (#SC-271509, Santa Cruz) at 4 °C on a rotating mixer overnight (16 h), followed by incubation with 30 µl pre-washed Protein G Agarose bead slurry (#11719416001, Roche) for an additional hour at 4 °C on a rotating mixer. For all IPs, beads were then washed four times with the respective IP wash buffer (50 mM Tris pH 7.5, 0.3% CHAPS; or 1% Triton X-100, 150 mM NaCl, 50 mM NaF) and boiled in 2x SDS loading buffer. Samples were analyzed by SDS-PAGE and the presence of co-immunoprecipitated proteins was detected by immunoblotting with appropriate specific antibodies.

#### *In vitro* kinase assay

*In vitro* ATM kinase assays were developed based on previous reports^67,69^, using FLAG-ATM immunopurified from HEK29FT cells treated with etoposide (20 μM, 2 h). In brief, cells of two near-confluent wells of a 6-well plate were lysed with modified TGN buffer (50 mM Tris pH 7.5, 150 mM NaCl, 1% Tween-20, 0.3% Nonidet P-40, 1 mM NaF, 1mM mM Na-vanadate), supplemented with 1x PhosSTOP phosphatase inhibitors (#04906837001, Roche) and 1x cOmplete protease inhibitors (#11697498001, Roche) for 10 min on ice. Samples were clarified by centrifugation (15000 rpm, 15 min, 4 °C), supernatants were subjected to immunoprecipitation by incubation with 30 μl pre-washed anti-FLAG M2 affinity gel (#A2220, Sigma) for 3 h (4 °C, rotating). Beads were then washed with TGN buffer with high salt (0.5 M NaCl), followed by two washes with kinase buffer (20 mM HEPES, pH 7.5, 50 mM NaCl, 10 mM MgCl_2_, 1 mM DTT, 10 mM MnCl_2_). Finally, the reactions were prepared by resuspending the beads in kinase buffer containing 125 μM unlabeled ATP, 6 μCi of [γ-^32^P] ATP (#SRP-301; Hartmann Analytic) and 1 μg of recombinant proteins as indicated in the figure. The kinase reaction was conducted at 30 °C for 1 h and stopped by addition of one volume 2x SDS sample buffer and boiling for 5 min at 95 °C. Samples were run on SDS-PAGE, the gel was dried (GD2000, Amersham) and radio-labeled proteins were visualized on PhosphorImager (Typhoon FLA9500, GE Healthcare).

#### Production of recombinant His_6_-tagged RHEB

Production of GST and GST-4E-BP1 were described in a previous report^7^. Recombinant His_6_-tagged RHEB protein was produced by transforming *E. coli* Rosetta electrocompetent bacteria with the pETM11-RHEB vector, according to standard procedures. In brief, protein expression was induced with isopropyl-β-D-thiogalactopyranoside (IPTG) for 4 h at 37 °C, and His_6_-RHEB was purified using Ni-NTA agarose (#1018244, Qiagen) and eluted with 333 mM imidazole (#A1073, Applichem).

#### Immunofluorescence and confocal microscopy

Immunofluorescence/confocal microscopy experiments were performed as described previously^64,68^. In brief, cells were seeded on glass coverslips coated with fibronectin (#A8350, Applichem) or poly-L-Lysine (sc-286689, Santa Cruz), treated as described in the figure legends, and fixed with 4% paraformaldehyde (PFA) (#28908, Thermo Scientific) in 1x PBS (10 min, room temperature), followed by a permeabilization step with 0.1% Triton in PBS for 10 min. Cells were blocked in blocking buffer (1% BSA in PBS) for 45 minutes. Staining with anti-TFEB (#4240, Cell Signaling Technology) and anti-γH2AX (#05-636, Sigma) antibodies (1:200 and 1:500, respectively in blocking buffer) was performed by incubation for 16 h at 4 °C. After staining with primary antibodies, coverslips were washed three times with PBS and then stained with highly cross-adsorbed fluorescent secondary antibodies (Donkey anti-Rabbit Alexa Fluor 488, Donkey anti-Mouse Alexa Fluor 594; both from Jackson ImmunoResearch) diluted 1:400 in blocking buffer for 1 hour at room temperature. Nuclei were stained with DAPI (#A1001, VWR) (1:2000 in PBS) for 10 min and coverslips were washed three times for 10 min with PBS before mounting on glass slides with Fluoromount-G (#00-4958-02, Invitrogen). All images were acquired on an SP8 Leica confocal microscope (TCS SP8 X or TCS SP8 DLS, Leica Microsystems) using a 63x oil objective lens. Image acquisition was performed using the LAS X software (Leica Microsystems). Images from single channels are shown in grayscale, whereas in merged images, Alexa Fluor 488 is shown in green and Alexa Fluor 594 in magenta. Brightness and contrast were adjusted for visualization purposes using Fiji (https://imagej.net/software/fiji/downloads)^70^. Alterations were applied to the entire image, keeping the parameters identical between all images of the same channel in each panel.

#### LysoTracker staining

For LysoTracker staining experiments, cells were seeded on fibronectin- or poly-L-lysine-coated coverslips and grown until they reached 80-90% confluency. Lysosomes were stained by the addition of 100 nM LysoTracker Red DND-99 (#L7528, Invitrogen) in the treatment media for 1 hour. Nuclei were stained by Hoechst 33342 (#1351304, Bio-Rad). Cells were then fixed with 4% PFA in PBS for 10 min at room temperature, washed and permeabilized with PBT solution (1x PBS, 0.1% Tween-20) for 10 min. Coverslips were mounted on slides using Fluoromount-G (#00-4958-02, Invitrogen). All images were captured on an SP8 Leica confocal microscope (TCS SP8 X or TCS SP8 DLS, Leica Microsystems) using a 63x oil objective lens. Image acquisition was performed using the LAS X software (Leica Microsystems). Brightness and contrast were adjusted for visualization purposes using Fiji^70^. Alterations were applied to the entire image, keeping the parameters identical between all images of the same channel.

#### Cell confluency measurements

Changes in cell confluency were measured using an IncuCyte S3 live-cell imaging and analysis System (Sartorius). In brief, cells were cultured in 12-well plates and images from nine different regions per well were acquired at regular intervals (8 h) with a 10x objective in an IncuCyte apparatus. Images were analyzed using the IncuCyte software.

### QUANTIFICATION AND STATISTICAL ANALYSIS

#### Statistical analysis

Statistical analysis and presentation of quantification data was performed using GraphPad Prism (versions 9 and 10). Data in all graphs are shown as mean ± SEM. For graphs with only two conditions shown (Fig. 1C,E,I and Fig. S1C), significance was calculated using Student’s t-test (unpaired, two-tailed). For all other graphs, significance for the indicated pairwise comparisons was calculated using one-way ANOVA with *post hoc* Tukey’s multiple comparisons test. Sample sizes (n) and significance values are indicated in figure legends (* p < 0.05, ** p < 0.01, *** p < 0.001, **** p < 0.0001, ns: non-significant).

All findings were reproducible over multiple independent experiments, within a reasonable degree of variability between replicates. For most experiments, at least three independent replicates were performed. The sample size for microscopy experiments (number of individual cells used in quantifications) is provided in the respective figure legends. No statistical method was used to predetermine sample size, which was determined in accordance with standard practices in the field. No data were excluded from the analyses. The experiments were not randomized, and the investigators were not blinded to allocation during experiments and outcome assessment.

#### Quantification of nuclear TFEB and LysoTracker intensities

Signal intensities were calculated using the Fiji software^70^. For nuclear TFEB quantification, nuclei were selected as regions-of-interest (ROIs) for approximately 50-60 cells over 4-5 independent representative images per condition acquired from one representative experiment (out of 3 independent replicate experiments) and integrated density was calculated, representing the sum of the values of all pixels in the given ROI. For quantification of LysoTracker signal, ROIs (which were individual cells) were determined for approximately 50-60 cells over 4-5 independent representative images per condition and integrated density was calculated, representing the sum of the values of all pixels in the given ROI.

## RESOURCE AVAILABILITY

### Lead Contact

Further information and reasonable requests for resources and reagents should be directed to and will be fulfilled by the Lead Contact, Dr. Constantinos Demetriades.

### Materials Availability

All unique plasmids and cell lines generated in this study are available from the corresponding author on reasonable request, with a completed material transfer agreement.

### Data and Code Availability

- The data that support the findings of this study (original immunoblot images and microscopy pictures) will be shared by the lead contact upon request.
- This paper does not report original code.
- Any additional information required to reanalyze the data reported in this paper is available from the lead contact upon request.

## Acknowledgements

We thank all members of the Demetriades lab for critical discussions; Julian Nüchel, Diana Terziyska, and Marija Kovacevic-Sarmiento for support with experiments; the MPI-AGE FACS & Imaging Core Facility for facilitating microscopy work; and Katrine Weischenfeldt for His-tagged RHEB protein purification. JP received support by the Cologne Graduate School of Ageing Research. CD is funded by the European Research Council (ERC) under the European Union’s Horizon 2020 research and innovation programme (grant agreement No 757729), and by the Max Planck Society. Parts of this work were supported by the ERC under the European Union’s Seventh Framework Programme via an ERC Starting Grant (grant agreement No 260602) to AAT; and by the Deutsche Forschungsgemeinschaft (DFG, German Research Foundation) through the Research Unit Grant FOR2722 (DE 3170/1-1; Project No 384170921) to CD. Graphical models in figures were created with BioRender.com.

## Author Contributions

Experimental work: J.P.; data analysis: J.P., C.D.; project design: J.P., A.A.T., C.D.; conceptualization: C.D.; supervision: C.D.; funding acquisition: A.A.T., C.D.; figure preparation: J.P., C.D.; manuscript draft: J.P., C.D. All authors approved the final version of the manuscript and agree on the content and conclusions.

## Declaration of Interests

The authors declare no competing interests.

## Notes

### Competing Interest Statement

The authors have declared no competing interest.

