## Supplementary material for "ATM-dependent RHEB phosphorylation couples DNA damage to lysosomal mTORC1 signaling to orchestrate the cellular response to genotoxic stress": Figures S1-S5

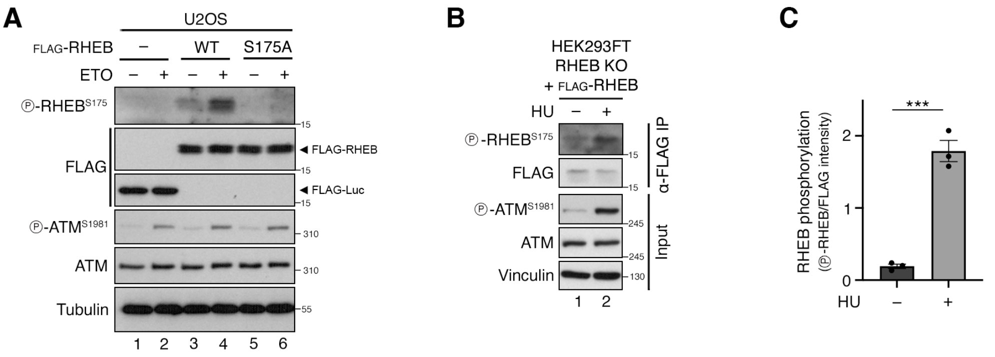


**Figure S1. Specific detection of RHEB$^{S175}$ phosphorylation is enhanced by genotoxic stress. Related to Figure 1.**

**(A)** Validation of the custom-made anti-phospho-RHEB$^{S175}$ antibody using a phospho-dead RHEB S175A mutant. Immunoblots with lysates from U2OS cells transiently expressing FLAG-tagged Luciferase (–), wild-type RHEB (WT), or RHEB$^{S175A}$ (S175A), treated with etoposide (ETO; 20 µM, 2 h) or DMSO as control and probed with different antibodies as indicated.

**(B-C)** Induction of genotoxic stress by treatment with hydroxyurea (HU) enhances RHEB$^{S175}$ phosphorylation. Immunoblots with anti-FLAG IP and input samples from HEK293FT RHEB KO cells stably expressing FLAG-tagged RHEB, treated with HU (1 mM, 2 h) or water as control (B). Quantification of RHEB phosphorylation (p-RHEB/FLAG signal) in (C). Data shown as mean ± SEM. *** $p < 0.001$. n = 3 independent experiments.

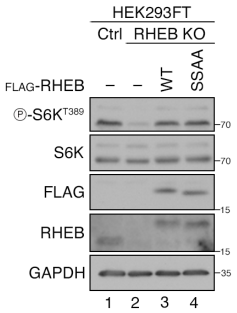


**Figure S2. Re-expression of RHEB$^{WT}$ or RHEB$^{SSAA}$ similarly restores mTORC1 activity in RHEB KO HEK293FT. Related to Figure 5.**

Immunoblots with lysates from HEK293FT control (Ctrl), RHEB knockout (RHEB KO), and RHEB KO cells stably expressing FLAG-tagged RHEB$^{WT}$ (WT) or RHEB$^{S6A/S175A}$ (SSAA), grown under standard culture conditions, probed with the indicated antibodies.

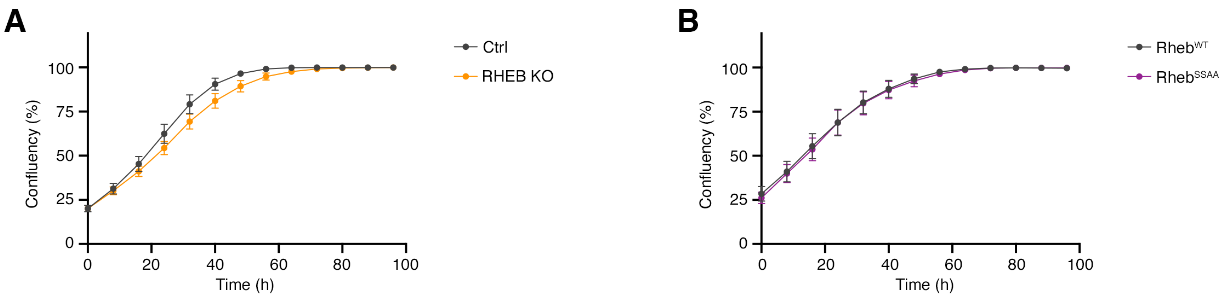


**Figure S3. Cells lacking RHEB or expressing a RHEB$^{SSAA}$ mutant display similar growth kinetics to control cells in the absence of DNA damage. Related to Figure 6.**

**(A)** Time-course of cell confluency (%) with control (Ctrl) and RHEB KO HEK293FT cells, grown under standard culture conditions.

**(B)** Time-course of cell confluency (%) with RHEB KO HEK293FT cells stably expressing FLAG-tagged wild-type RHEB (WT) or a phosphorylation-deficient RHEB$^{S6A/S175A}$ mutant (SSAA), grown under standard culture conditions.

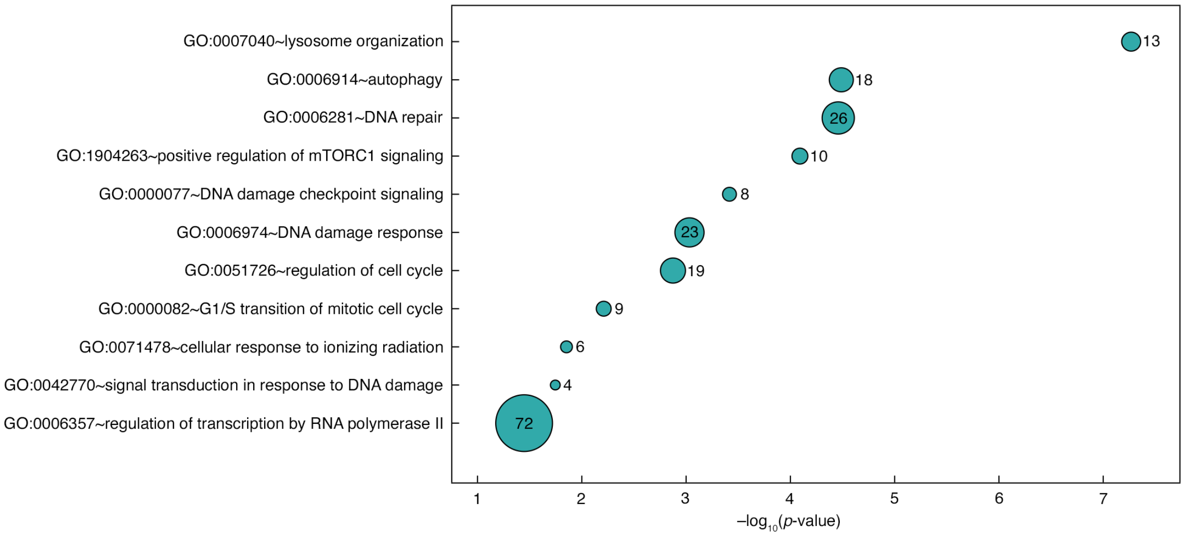


**Figure S4. Biological Process (BP) GO term analysis of TFEB target genes reveals enrichment of DDR-related pathways. Related to Figure 6.**

Dot plot showing significantly enriched BP GO terms among curated TFEB target genes. Circle size reflects the number of TFEB target genes associated with each term.

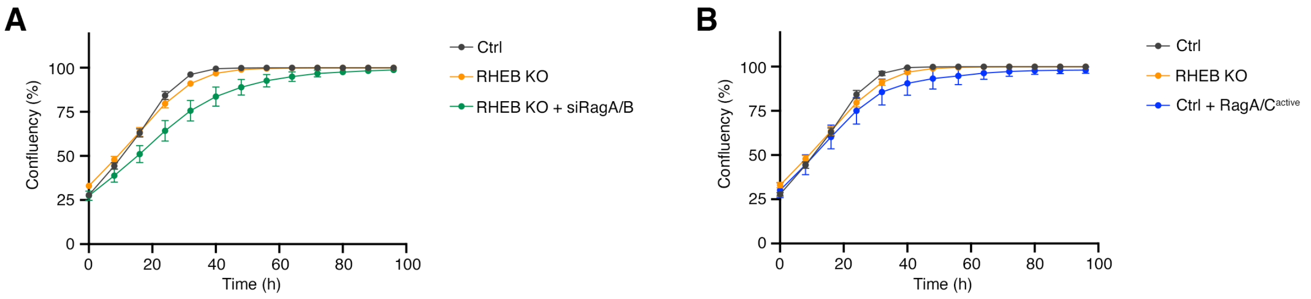


**Figure S5. Growth kinetics of control and RHEB KO cells upon expression of active RagA/C dimers or RagA/B knockdown in the absence of DNA damage. Related to Figure 7.**

**(A)** Time-course of HEK293FT cell confluency (%) with controls (Ctrl), RHEB knockouts (RHEB KO), or RHEB KO cells with RagA/B knockdown (RHEB KO + siRagA/B), grown under standard culture conditions.

**(B)** Time-course of HEK293FT cell confluency (%) with controls (Ctrl), RHEB knockouts (RHEB KO), or cells transiently expressing active RagA/C dimers (Ctrl + RagA/$C^{active}$), grown under normal cell culture conditions.
